# Parallel adaptation to higher temperatures in divergent clades of the nematode *Pristionchus pacificus*

**DOI:** 10.1101/096727

**Authors:** Mark Leaver, Merve Kayhan, Angela McGaughran, Christian Roedelsperger, Anthony A. Hyman, Ralf J. Sommer

## Abstract

Studying the effect of temperature on fertility is particularly important in the light of ongoing climate change. We need to know if organisms can adapt to higher temperatures and, if so, what are the evolutionary mechanisms behind such adaptation. Such studies have been hampered by the lack different populations of sufficient sizes with which to relate the phenotype of temperature tolerance to the underlying genotypes. Here, we examined temperature adaptation in populations of the nematode *Pristionchus pacificus*, in which individual strains are able to successfully reproduce at 30°C. Analysis of the frequency of heat tolerant strains in different temperature zones on La Réunion supports that this trait is subject to natural selection. Reconstruction of ancestral states along the phylogeny of highly differentiated *P. pacificus* clades suggests that heat tolerance evolved multiple times independently. This is further supported by genome wide association studies showing that heat tolerance is a polygenic trait and that different loci are used by individual *P. pacificus* clades to develop heat tolerance. More precisely, analysis of allele frequencies indicated that most genetic markers that are associated with heat tolerance are only polymorphic in individual clades. While in some *P. pacificus* clades, parallel evolution of heat tolerance can be explained by ancestral polymorphism or by gene flow across clades, we observe at least one clearly distinct and independent scenario where heat tolerance emerged by *de novo* mutation. Thus, temperature tolerance evolved at least two times independently in the evolutionary history of this species. Our data suggest that studies of wild populations of *P. pacificus* will reveal distinct cellular mechanisms driving temperature adaptation.

## Background

A fundamental question in biology is what are the evolutionary mechanisms that drive the adaptation of organisms to the environmental conditions of their niche. Studying the evolution of natural populations has shown that common solutions are adopted during adaptation to similar niches. For example, populations of threespine sticklebacks have been shown to independently acquire the same phenotype during repeated adaptation to fresh-water habitats [1,2]. Similarly, during the adaptive radiation of cichlid fish in the lakes of East Africa, there have been examples of parallel [3] and convergent evolution, giving rise to striking examples of species adopting similar body-forms while adapting to the same ecological niche [4]. The environment, therefore, can be a selective factor that drives the evolution of new species [5].

An important environmental variable is temperature, which fluctuates both on daily and yearly cycles. Although some information is available about how temperature affects fitness, especially in ectotherms (cold-blooded organism) [6–8], little is known about if natural populations adapt to temperature. Furthermore, little is known about the evolutionary mechanisms behind their adaptation or about the genes that influence it. Studying adaptation to temperature is particularly important in the light of increasing evidence for climate change [9].

Nematodes, such as *Caenorhabditis elegans*, are excellent model organisms to study the effect of temperature on the fitness of ectotherms. Temperature is known to affect their fertility [10,11] and that of its sister species, *C. briggsae* [12]. There is also evidence that *C. elegans* and *C. briggsae* have adapted to different temperature niches, with *C. elegans* being more prevalent in temperate regions and *C. briggsae* being more prevalent in the tropics [12]. However, to understand the dynamics of this adaptation and its genetic basis, we need to study this in wild isolates of the same species. Another nematode that has been developed as a model organism for studying evolutionary biology is *Pristionchus pacificus* [13]. *P. pacificus* is associated with beetles, with which it has a necromenic relationship. Specifically, it can be found on beetles in a dormant state called the dauer stage [14,15] and, once the beetle has died, nematodes re-enter the reproductive cycle and feed on the bacteria growing on the decomposing body of the beetle [16]. *P. pacificus* is associated with scarab beetles and stag beetles, and has undergone host switching to utilize different beetle hosts found in novel habitats. This association has enabled the collection of large numbers of natural isolates of *P. pacificus* [14]. As of today, more than 1000 wild isolates of *P. pacificus* have been obtained from around the world [17].

The Indian Ocean island of La Réunion is particularly important for understanding adaptation of *P. pacificus* to different temperature niches, because the island experiences a range of temperatures to which diverse populations of nematodes could have adapted. La Réunion has been colonized many times independently by representatives of all major clades of *P. pacificus* [14,18,19]. These clades of *P. pacificus* (formerly designated A-D) are distinguished by mitochondrial markers [18]. However, analysis of whole genome sequencing data has revealed that these mitochondrial clades actually represent at least six genome-wide clades [20], of which four are predominantly found on La Réunion (A2, B, C, D). The colonization by clade C of La Réunion was most likely achieved via the ancestor of the scarab beetle *Oryctes borbonicus*. Subsequently, clade C strains have spread across most of the western part of the island [14]. *O. borbonicus* is endemic to La Réunion as is *P. pacificus* clade C, suggesting a long history of co-evolution between nematode and beetle [21]. Other beetle species have been introduced to the island more recently. For example, *Maladera afftnis* was introduced between 800-1800 [22] and is currently host to clade A strains [14]. Competition between recently arrived and well-established beetle hosts has presumably limited nematode dispersal, leading to distinct, occasionally overlapping populations distributed across the island [23]. Clade B strains are unique to the island and form an isolated population at high altitudes, where they are found in association with the beetle *Amneidus godefroyi* [14,24]. Since colonizing the island, individual clades of *P. pacificus* continue to accumulate genetic and phenotypic variations as they adapt to local conditions [23]. This has led to a high degree of heterogeneity and population structure, particularly in clade C [18]. While strains of different clades differ in about 1% of their sequence, strains of the same clade exhibit lower levels of nucleotide diversity (about 0.1%) comparable to the global diversity of *C. elegans* [20]. Similarly, while recombination frequently occurs within clades, admixture between different clades is rarely observed [18]. Together with evidence for intraspecific competition, this suggests a process of incipient speciation in *P. pacificus* [25,26].

We have previously shown that high temperature differentially affects fertility among *P. pacificus* natural isolates [27]. An isolate from California can no longer give rise to fertile offspring at 30°C, while one from Japan remains fertile at that temperature. The phenotype of the Japanese strain is designated as high temperature tolerant or Htt. We have shown that a single locus on chromosome V in the Japanese strain is responsible for the Htt phenotype and that there is some variation in the phenotype among natural isolates from world-wide locations [27]. However, it is not yet known if this phenotype is subject to natural selection in wild populations. As a volcanic island, La Réunion encompasses a range of ecotypes formed by its diverse geology, biodiversity and weather [23]. More importantly, the island ranges from warm coastal regions that experience temperatures up to 36°C to the cooler slopes of the volcano that experience temperatures up to 22°C. This model system offers the perfect opportunity to test if the Htt phenotype is subject to natural selection in populations of nematodes and, if so, to investigate the evolutionary mechanisms behind such adaptation. Furthermore, we can combine this evolutionary approach with a genetic one to investigate which loci are associated with temperature adaptation.

Here, we assay the temperature tolerance of 289 strains of *P. pacificus*, collected from different sites on the island experiencing a range of temperatures. We show that temperature acts as a selective factor on natural populations of nematodes and further show that high temperature adaptation has evolved at least twice in an example of parallel evolution.

## Results

### Populations of nematodes on La Réunion are subject to natural selection

We define the Htt phenotype as the ability to give rise to fertile offspring at 30°C. To test if this phenotype is subject to natural selection, we investigated the occurrence of the Htt phenotype in a large collection of natural isolates from La Réunion. A simple test for this is to look for a correlation between the frequency of the trait and the selective factor [28], in this case altitude as a proxy for temperature (Sup. Fig. 1A). We therefore predicted that strains that were isolated at higher altitudes (lower temperatures) should be less likely to be Htt than strains that were collected at lower altitude locations (higher temperatures). To test this, we utilized the existing collection of natural isolates of *P. pacificus* from La Réunion, collected between 2008 and 2011, that are known to represent different genetic populations [18,25]. As there were relatively few strains collected from below 500 m because of sheer cliffs, urban areas and sugar cane farming at lower altitudes, we identified new locations with relatively un-spoilt habitat and collected additional strains between 2012 and 2015 (Sup. Fig. 1B and see methods). Collecting at these sites gave enough *P. pacificus* isolates to complete sampling from sea level (covering a range of maximum daily temperatures from 22-36°C). These strains cover the diversity of genomic clades that are found on the island [25] and are from geographically separated populations that may have adapted to local conditions.

For phenotyping, individual J3 larvae maintained at 20°C were shifted to 30°C then, after seven days of incubation, worms were inspected for their ability to give rise to fertile offspring. Strains were scored either as Htt if they gave fertile offspring or as Ltt if they were infertile or only gave rise to infertile offspring (see methods for details). In total, 21.5% of the 289 strains were Htt (Sup. Table 1). As predicted, low altitude locations tended to have more Htt strains, for example, 76.7% (n=30) of strains from Saint Benoit (SB, 22 m altitude) were Htt (Sup. Table 1). In contrast, no strains from locations above 1000 m were Htt (n=73). To test for a correlation, the occurrence of the Htt phenotype was plotted by altitude (Fig. 1). The percentage of strains giving rise to fertile offspring at 30°C was as high as 77% at the lowest altitude bin and only 6.9% for the bin centered at 1075 m (Fig. 1). A subset of these data fell onto a straight line (R^2^=0.730), showing a correlation between altitude and high temperature tolerance. Analysis of variance (Anova, see methods) showed that the correlation between the occurrence of the phenotype and the altitude at which any strain was collected was highly significant (p-value=2.2×10^−16^). These findings show that the occurrence of the Htt phenotype is related to environmental temperature and is therefore likely to be subject to natural selection.

**Figure 1.**
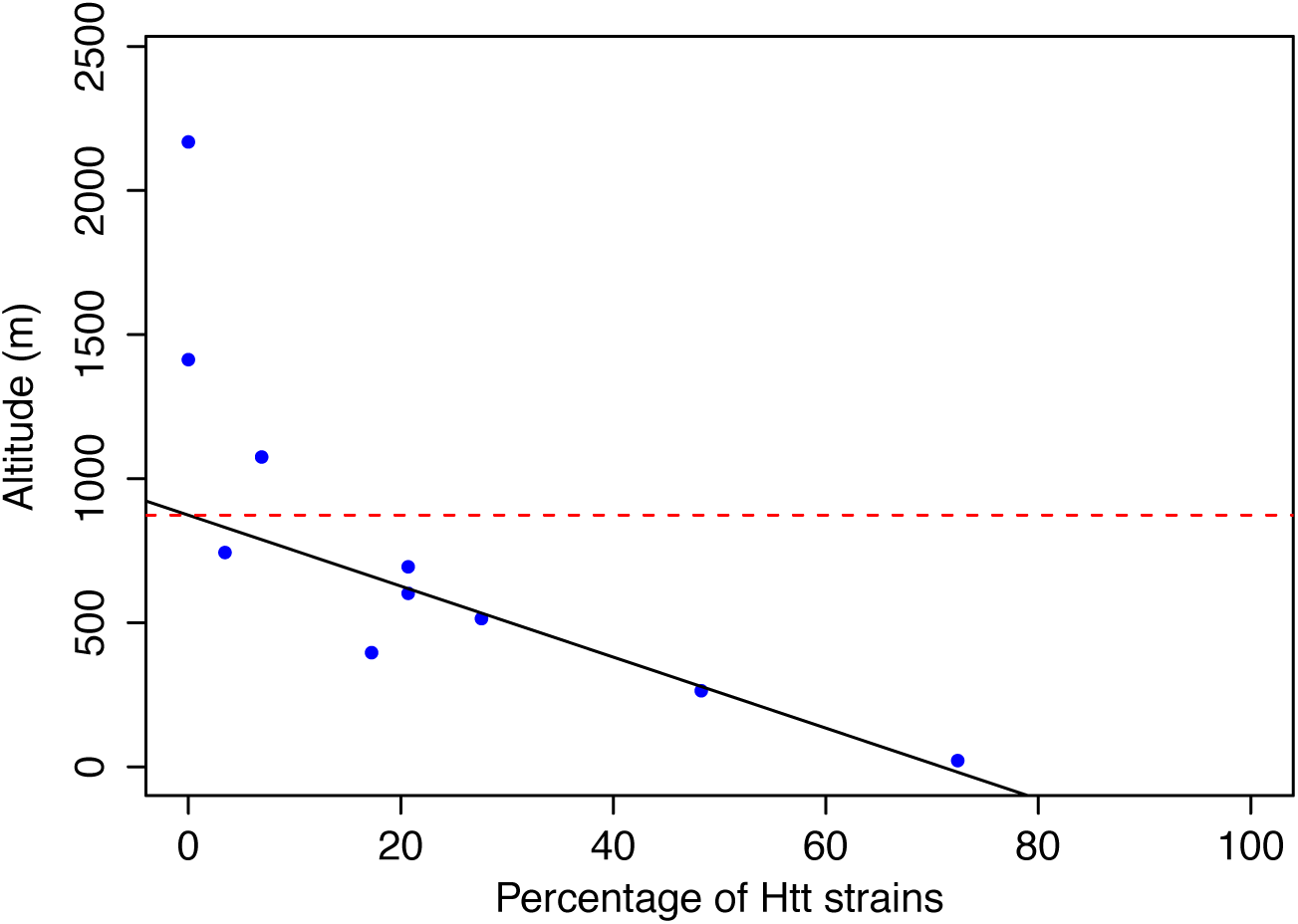
The occurrence of the Htt strains decreases with increasing altitude of their collection site. The correlation between altitude and the occurrence of Htt strains. Strains were ranked by the altitude of their collection site then grouped into 10 bins. The percentage of high temperature tolerant strains in each bin was calculated and plotted against the average altitude of the sample site of all strains in that bin. The black line is a fit for the data between 0 and 1000 m. The red line is the altitude (867 m) where the percentage of Htt strains tends to zero as extrapolated from the fit.

### Strains isolated from beetles found at lower altitudes are more likely to be Htt

The beetle species from which nematodes were isolated have different habitat ranges corresponding to their preferred altitudes, as quantified in Sup. Fig. 2. For example, *A. godefroyi* is found only at higher altitudes [24] while other species, including *M. affinis*, prefer lower altitudes [22]. The occurrence of Htt strains isolated from different beetle species should differ depending on the temperatures of the beetle’s habitat. Indeed, Htt strains were found more frequently on *M. affinis* and *Aphodius sublividus* (found at low altitudes), but were not found at all on *A. godefroyi* and *O. borbonicus* (found at higher altitudes, inset Fig. 2). Anova also showed that the percentage of Htt strains found on different species of beetles deviated from a random pattern (p-value=2.5×10^−11^).

**Figure 2.**
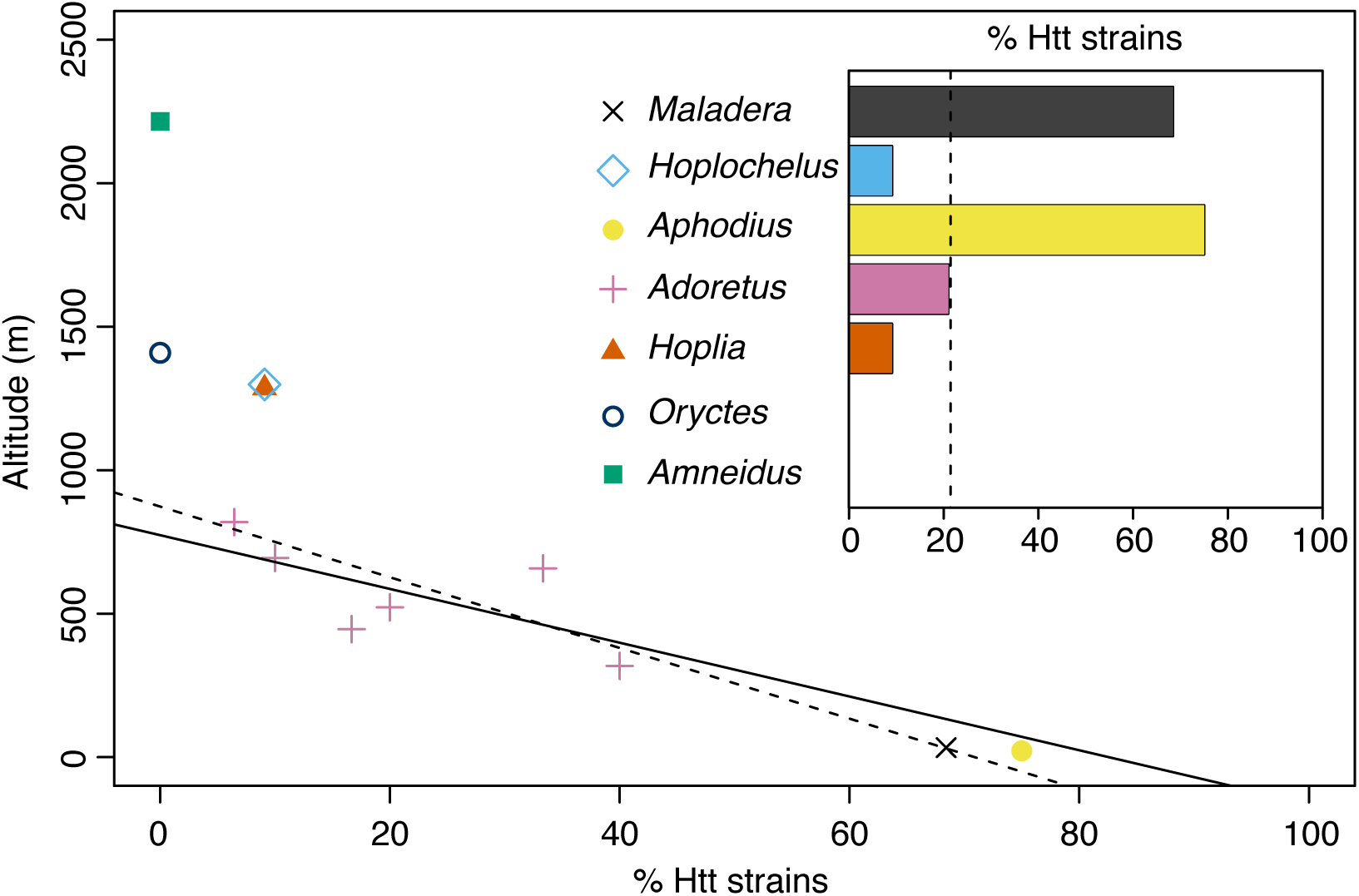
The correlation between the percentage of Htt strains and altitude remains after accounting for the beetle host. Inset: the percentage of nematodes that are high temperature tolerant for strains isolated from each beetle species. Main panel, the percentage of Htt strains isolated from each beetle plotted against the mean altitude of their collection site. For nematodes isolated from *Adoretus sp*., strains are grouped into bins and each bin plotted separately and fit with a straight line (dashed line). The solid line is the fit for the data as plotted in Fig.1.

To eliminate any influence that different beetle hosts may have on the occurrence of Htt strains, we plotted the data of strains isolated from different beetles separately (Fig. 2). The largest number of nematode strains was isolated from *Adoretus sp*. (n=181/289), which in contrast to other beetles, is found at a range of altitudes. Indeed, *Adoretus*-derived strains show a linear correlation between altitude and percentage of Htt strains and the correlation between altitude and percentage of Htt strains remains, although weaker than that seen in the whole dataset (R^2^=0.457, compare Figures 1 and 2). From this it can be concluded that different beetle species carry different frequencies of Htt strains, and that, in the one beetle species found at a range of altitudes (thus, for which the effect of beetle can be factored out), the correlation between altitude and Htt remains.

### Strains that belong to clade A are more likely to be Htt

Next, we considered the evolutionary relationship between strains of *P. pacificus* as another potential influencing factor on the frequency of Htt strains. For example, strains belonging to clade B might respond differently to temperature because they form an isolated population [24] that might be susceptible to the founder effect [29]. Therefore, the percentage of Htt strains belonging to each of the four mitochondrial clades was calculated. Figure 3 shows that clades found at high altitudes have no Htt strains. In contrast, clade A strains, which are more prevalent at lower altitudes, have a higher percentage of Htt strains. Anova showed that the percentage of high temperature tolerant strains found in different clades deviated from that which would be expected purely by chance (p-value = 6.6×10^−14^). From this it can be concluded that a strain will be more or less likely to be Htt depending on which clade it belongs to.

**Figure 3.**
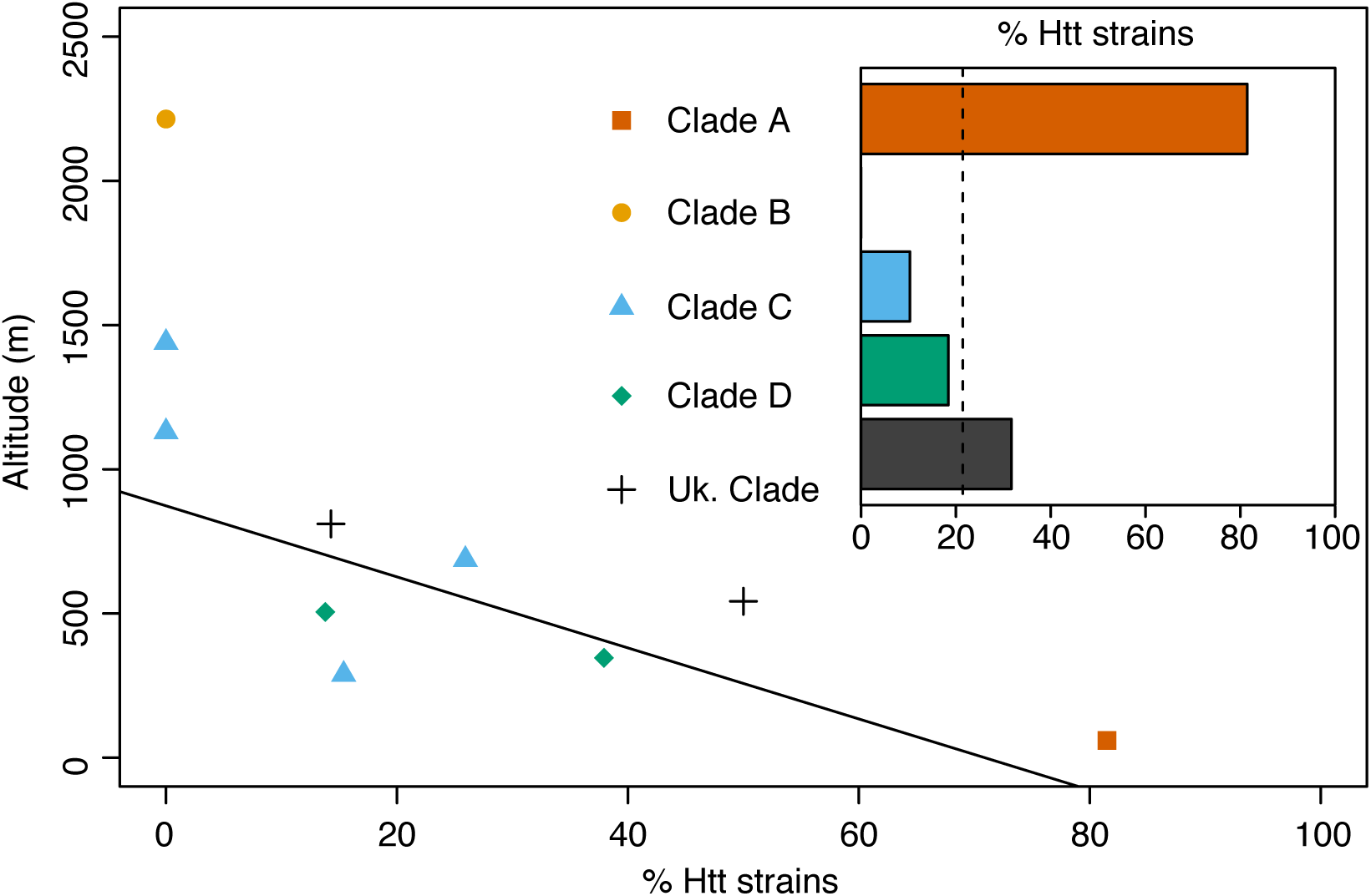
The occurrence of Htt phenotype in strains that belong to different clades. Inset: the percentage of strains that are Htt belonging to each clade. Main panel: the percentage of Htt strains belonging to each clade is plotted against the mean altitude of their collection site. Mitochondrial clades A, B, C, D and strains of unknown clade were grouped into bins and each bin plotted separately.

Of the three influencing factors (beetle, clade and altitude), the influence of beetle can be eliminated by analysing strains isolated from *Adoretus sp*. (Fig. 2), but the influence of clade and altitude remain to be disentangled. In an attempt to distinguish which variable best explained the occurrence of the trait, we performed model selection analysis using Bayesian Information Criterion (BIC). BIC is a statistical tool that selects from a set of models the one that best explains the observed data using a likelihood function [30]. The BIC analysis showed that altitude explained the variation in frequency of the Htt trait better than beetle or clade (0.99 for altitude, 8.7×10^−11^ for clade and 3.0×10^−6^ for beetle: the largest value indicates the variable that best fits the data, see methods for details). This supports our assertion that temperature is likely to be the main influencing factor on the phenotype, as expected from our phenotype assay.

### Phylogeny of world-wide strains of P. pacificus suggests parallel evolution of the Htt phenotype

To investigate the evolution of temperature tolerance in a larger collection of *P. pacificus* strains, we combined the current dataset with strains from world-wide locations that we had analyzed previously [27]. SNPs were identified by comparing whole genome data of these strains. These SNPs were then concatenated and the sequence used to construct a phylogenetic tree by the maximum likelihood method with ultrafast bootstrapping [31,32] (Fig. 4). The world-wide diversity of *P. pacificus* consists of six major clades based on genome-wide markers; clades A1, A2, A3, B, C and D, as well as one divergent strain used to root the tree (Fig. 4), consistent with previous work [20,27,33]. Clades C and D consist largely of strains from locations spread all over La Réunion and nearby island Mauritius; clade B consists exclusively of strains found on La Réunion at locations higher than 2100 m; and clades A1, A2 and A3 consist of strains from worldwide locations, La Réunion and Mauritius. Clade A3 includes the wild type reference strain, RS2333.

**Figure 4.**
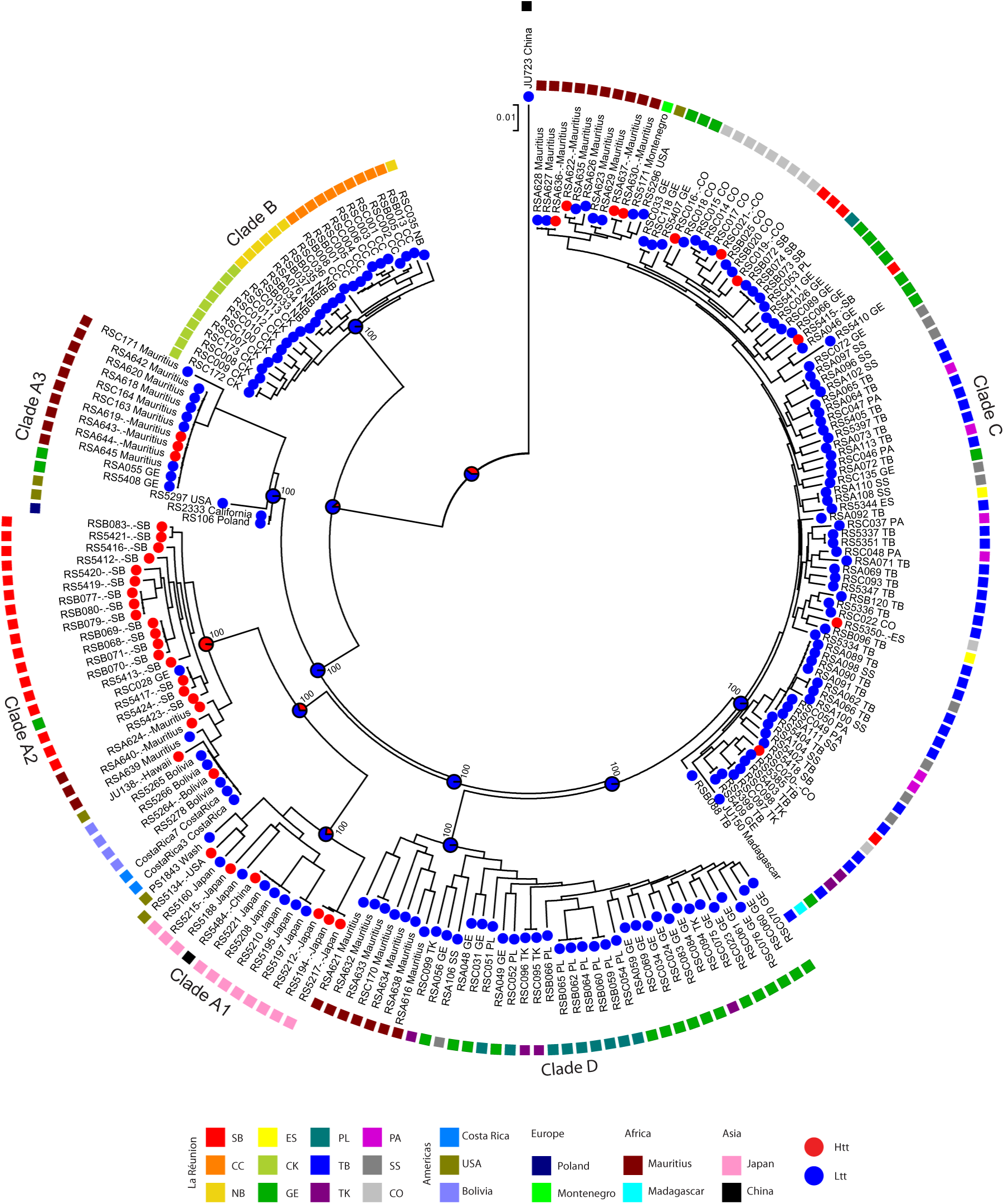
Phylogenetic tree of *P. pacificus* strains shows one clade highly enriched with Htt strains. A maximum likelihood tree based on the sequence of 873932 concatenated SNPs across the whole genome for 212 strains. The phenotype of each strain is indicated by a circle at the terminal node of the tree (red = Htt, blue = Ltt). The sample site is indicated by a coloured square. For the nodes of the major clades, the degree of bootstrap support and the inferred phenotype is indicated by pie charts at the internal nodes.

Htt strains are sporadically scattered throughout clade C (Fig. 4) suggesting that there were recent and independent acquisitions of high temperature tolerance in strains from La Réunion and Mauritius [27]. There are also Htt strains in clades A1, A2 and A3, including sub-clades with many Htt strains or single strains from diverse locations (Fig. 4). To test if temperature tolerance has arisen multiple times independently, we inferred the phenotype of ancestral strains using a Bayesian algorithm (Fig. 4 and Sup. Fig. 3). This analysis predicts that the last ancestor of all *P. pacificus* strains was likely to be low temperature tolerant. Furthermore, the ancestor of deeply rooted clade B is predicted to have been low temperature tolerant and the same is true for clades C, D and A1 (Fig. 4). In contrast the ancestor of clade A2 is predicted to have been high temperature tolerant (Fig. 4 and Sup. Fig. 3). This suggests that the Htt phenotype is a recent and derived state and could have arisen *de novo* within clades A1, A2, A3 and C, consistent with parallel evolution. The alternative explanation is that the Htt phenotype arose once and was transferred between clades by recombination during outcrossing.

### Reversion of strains back to the Ltt phenotype from an Htt ancestor

A closer examination of strains from clade A2 shows that there are many closely related strains that are Htt and form a “hot” sub-clade (Figs. 4). These strains were collected from SB on La Réunion, which has an altitude of 22 m and experiences a maximum annual temperature of 34°C. Interestingly, there is one example of a reversion in this sub-clade: strain RSC028, which is Ltt, was collected at Grand Etang (GE, 527 m), where temperatures are lower than at SB. There are further examples of reversion in clade A2: for example, the event that gives rise to a “cold” sub-clade (Fig. 4 and Sup. Fig. 3). Interestingly, two strains in this sub-clade, JU138 and RS5264, have regained the Htt phenotype. From this, we can conclude that there are examples of reversion of the Htt trait back to the ancestral state as strains adapt to local conditions.

### Htt is a polygenic trait

To better understand the evolution of the Htt phenotype, we wanted to identify SNPs that are associated with the phenotype so that we could later test if their pattern of inheritance indicated that they had arisen by *de novo* mutation or by recombination between clades. Genome wide association studies (GWASs) of model organisms have been used to identify SNPs linked to ecologically important phenotypes [34], including pH tolerance in *P. pacificus* [25]. Here, we performed GWAS using the easyGWAS online platform [35] and selected 223 strains for which whole genome sequence and temperature tolerance data are available. These strains were a subset of the La Réunion strains as well as strains from world-wide locations. GWAS was performed using the EMMAX algorithm to account for population structure and a minor allele frequency cut-off of 10%. The algorithm identified 32117 SNPs after filtering and inspection of the Q-Q plots revealed evidence for population stratification (genomic control, *λ*=0.55), which could be eliminated by accounting for the beetle host as a covariate and removing highly divergent strains from the analysis (*λ*=0.99), although this greatly reduced the number of SNPs (4447). Together, the EMMAX algorithm and the covariate take into account the two influencing factors we identified as affecting the occurrence of the trait. The hits from our GWAS were scattered over all six chromosomes with a total of 12 SNPs that were significantly associated with the Htt phenotype (Sup. Table 2). These SNPs are either in genes that affect the phenotype or are linked to the causative mutation. This finding suggests that SNPs in multiple genes influence temperature tolerance, i.e. that Htt is a polygenic trait.

### Clade-specific GWAS and allele-frequency analysis support independent gains

A study of the genetic causes of hypertension [36] showed that polymorphisms linked to the disease could be better identified by studying individuals in a subpopulation where there was an increased occurrence of the condition. The equivalent for our study would be to independently analyse clades A1 and A2 combined because the percentage of Htt strains is higher in these clades than in the complete dataset. In comparison, we considered clade C independently because the occurrence of Htt strains is lower here. GWAS analysis of these strains revealed that these subpopulations also had low levels of stratification (with *λ*=1.00 and *λ*=1.16 for clade A1+A2 and clade C respectively). GWAS of clade A1 and A2 strains gave one cluster of a few SNPs on chromosome V including one that fell above the significance threshold (Sup. Table 2 and results online). In contrast, GWAS of clade C showed a broad region of chromosome I with several SNPs with similar –log(p-level) that also fell above the threshold. The finding that the different GWASs identify loci on different parts of the genome is consistent with independent acquisitions of the phenotype in clade C and clades A1/A2.

To further test whether different loci have been used by individual clades to evolve heat tolerance, we investigated patterns of fixation of candidate SNPs that were identified by GWAS. If heat tolerance evolved independently in each clade, then the SNPs that are associated with the phenotype should have alleles that are unique to one clade and not polymorphic in other clades. Indeed, most of the identified SNPs are polymorphic in just a single clade (highlighted Sup. Table 3). For example, ChrI:19341896 is polymorphic in clade C (either a T or an A) but always an A in all other clades. This observation and other clade C SNPs (highlighted in blue in Sup. Table 3) supports a *de novo* mutations specific to clade C. This suggesting that there was an independent acquisition of Htt in clade C, and another in clades A1, A2, or A3. Alternatively, if heat tolerance was a polygenic trait that evolved once, signals that are captured by the GWAS should come from SNPs that are polymorphic in all clades exhibiting heat tolerant strains. Indeed, we do see examples of SNPs that are polymorphic in clades with Htt strains (highlighted in yellow in Table 3), for example ChrV:13634723 is polymorphic in clade A1 and clade C (either a T or a C) but does not vary among strains from the other clades. This suggests that recombination might transfer Htt alleles between clades. We also observe examples of SNPs that are polymorphic in one clade, with both alleles found in other clades that are not polymorphic. For example, ChrI:18417120 is polymorphic in clade A1 (either a G or an A) but not in other clades where it is always either a G or an A. The most likely explanation for this is that such positions were polymorphic in the ancestral population, selected for in Htt strains (in clade A1 in the example), and differentially fixed in the other clades. Together, these findings suggest that heat tolerance evolved separately in some clades by *de novo* mutation and/or evolved in other clades by fixation of ancestral polymorphisms and subsequent spread by recombination between clades.

## Discussion

Because of ongoing climate change, there is increased interest in the extent and speed by which natural populations can adapt to higher temperatures and the evolutionary mechanisms by which they do so. Here, by comparing the temperature tolerance of 289 strains from La Réunion, we show that natural populations of *P. pacificus* are indeed subject to natural selection and have gained the ability to remain fertile at 30°C. Because we have estimates of the number of generations it took to generate the diversity of each of the major clades [37,38], we can estimate the amount of time it took for this adaptation to occur. For instance, in clade C, within 5.6×10^5^ generations, a low temperature tolerant strain had adapted by 2°C to a high temperature tolerant strain. A generation is likely to range between 3 days and 3 months, giving a time frame for adaptation between approximately 4600 and 140000 years (see methods).

Phylogenetic analysis of *P. pacificus* strains suggested that the Htt phenotype has evolved multiple times independently in the different clades in an example of parallel evolution. Alternatively, Htt could have evolved once and either remained polymorphic during the divergence between clades, or transferred between clades by recombination. However, two previous results suggest that this scenario would occur only rarely. First, analysis of mutation accumulation lines indicates that different clades are separated by millions of generations and it seems unlikely that polymorphisms would be maintained for such long periods [38]. In accordance with this, comparison of nucleotide diversity between clades shows very little (<7%) shared variation [20]. Second, analysis of linkage disequilibrium and population structure shows that, while there is frequent recombination within clades, admixture between different clades seems to be rare [18,20]. In summary, the analysis of the evolution of Htt along the phylogeny of *P. pacificus* is compatible with a scenario where Htt has evolved multiple times independently.

We also show examples of reversion back to low temperature tolerance, which occurs most likely by recombination within clades. The presence of reversions suggests that selection is not strong enough to ensure fixation of the phenotype. Alternatively, selection can occur in the other direction, i.e., there might be some balancing selection acting against fixation. Documented examples of reversion in natural populations are rare and could only be observed in this case because of the large collection of phenotyped strains and a detailed and accurate phylogeny.

GWAS identified SNPs associated with the Htt phenotype that were scattered across the chromosome, suggesting that different mutations can lead to the same phenotype, possibly by affecting different genes in the same pathway that regulates high temperature tolerance. The GWAS of all strains in our study gave 12 SNPs that were significantly associated with the phenotype. This represents SNPs from the worldwide diversity of *P. pacificus* that are either in genes or regulatory regions that directly affect the phenotype, or that are linked to the causative mutation. The GWAS of strains from clades A1 and A2 gave only one SNP associated with the phenotype. This SNP, intragenic between *gst-24* (glutathione S-transferase) and *cyp-29A4* (cytochrome P450, Sup. Table 4) was also found in the GWAS of all clades, showing that the large number of Htt strains in clade A could give a signal strong enough to be statistically significant in the whole dataset. Interestingly, this SNP is on chromosome V and is close to the region that we identified in pair-wise mapping of the Htt phenotype in a clade A1 Japanese strain (RS5194) [27]. The GWAS of clade C strains gave SNPs with significant scores mostly on chromosome I, but also on II and V. The 23 hits on chromosome I have the same or similar –log(p-value) and form a cluster of SNPs adjacent to each other. This resembles a linkage group and might represent an example of hitchhiking, where one or a few SNPs which cause the phenotype are selected for and carry with them the linked SNPs by recombination [39]. Such a scenario would be expected for a trait that is passed between individuals by sexual reproduction. The observation that GWAS of strains from different clades identified different SNPs is again consistent with parallel evolution of the phenotype in clade C and clades A1/A2.

In an attempt to further investigate the evolution of the Htt phenotype, we measured allele frequencies for the GWAS hits. This showed that most of the identified SNPs were polymorphic in just a single clade, which is consistent with different loci being employed by individual *P. pacificus* clades to become high temperature tolerant. Alternatively, in theory an ancestral population that was polymorphic at all Htt loci could have evolved once. Subsequently, Htt could have been lost in individual clades by fixation or could have been regained by recombination across clades. This alternative scenario is consistent with the finding that for most significant SNPs, both alleles are fixed in at least one other clade. However, even if this alternative scenario holds true for most Htt loci, we observe one clearly distinct and therefore independent pattern in clade C. All SNPs on chromosome I with significant association to Htt in the clade C specific GWAS are polymorphic only in clade C and only one of the alleles is present in all other clades. This indicates that clade C specific alleles in this region represent recently derived alleles that evolved *de novo* and therefore could not have been transferred by recombination across clades. Thus, we conclude that Htt evolved at least two times independently during the evolutionary history of *P. pacificus*.

Based on our findings and existing information about populations of *P. pacificus* on La Réunion, we can suggest a plausible series of events leading up to the current distribution of strains on the island. The common ancestor of clade C was most likely low temperature tolerant and, during diversification on the island, strains from that clade have gained the Htt phenotype at locations where temperatures are higher (SB, ES and CO, Fig. 4). The opposite seems to the case for clade A, whose ancestral population had most likely already evolved Htt (Fig. 4 and sup. Fig. 3) making it ideally suited to fill the high temperature niche that is now SB, forming the hot sub-clade in clade A2. Strong selective pressure at this location meant that descendants of these strains kept the ability to survive at high temperatures. The ancestor of strain RSC028 was part of this Htt population at SB and was likely transported to GE (527 m) where temperatures are lower, was subjected to lower selective pressure or balancing selection, and then reverted to Ltt.

The fact that there are many SNPs associated with the phenotype suggests that the phenotype is polygenic with many loci influencing the phenotype. We therefore conclude that temperature tolerance is a complex trait, possibly a threshold trait. Thresholds traits have an underlying continuous nature with a threshold above which the trait manifests itself [40]. The Htt phenotype fits this definition because each strain could have an upper temperature limit that falls onto a continuous scale and our definition of Htt involves a threshold of 30°C. Future work could confirm this by precisely measuring the upper temperature limit of strains which have upper temperature limits between those of Htt and Ltt strains. Adaptation to temperature has previously been reported for *Arabidopsis thaliana* [41] and *Drosophila melanogaster* [42]. There have also been several attempts to map loci responsible for the temperature dependence of fertility related traits in *C. elegans* [43–46]. These studies could only identify large genomic regions containing many candidate genes because of the low marker densities used. Also, these studies utilized recombinant inbred lines made between lab adapted strains of *C. elegans*, which would mean that the ecological relevance of these phenotypes is limited [47].

We do not yet know the underlying cell biology mechanisms behind temperature adaptation. However, the ease by which the cell biology can be studied in these organisms suggests that it will be possible to directly relate adaptation to a temperature niche to the underlying cell biology mechanisms in the near future.

## Methods

### Collection of natural isolates of P. pacificus

Strains of *P. pacificus* had previously been collected by the Sommer lab (prefix RS), the Sternberg lab (prefix PS) and Félix lab (JU) from world-wide locations including many from La Réunion. To increase the number of strains for analysis, more strains were collected from La Réunion. See [16] for details of collection methods. Beetles were dissected along the midline and placed onto NGM plates seeded with *Escherichia coli* OP50. After three days of incubation, plates were inspected for gravid hermaphrodites using a dissecting microscope. The SSU gene was sequenced to confirm strains as *P. pacificus*.

### Selection of sampling sites for temperature transect

Preliminary analysis set of 149 strains collected from La Réunion by the Sommer lab from 2008-2010 showed that one location had a high percentage of Htt strains. However, this dataset was lacking sufficient locations from 28-700m to properly test for a correlation between altitude and the occurrence of Htt strains. We identified locations around Plane d’ Affourches (PD5-PD7) and Saint Phillipe (SP) where *Adoretus sp*. could be found. Sampling over the subsequent years (2011-2015) collected enough beetles to give nematodes from locations covering the full range of altitudes.

### Strain maintenance

Worms were propagated on 0P50/NGM plates according to standard methods for *C. elegans* [48]. In order to minimise epigenetic effects, all worms were kept at 20°C for at least two generations on abundant food before temperature experiments were performed. The incubators used for routine maintenance were Heraeus BK 6160 incubators (Thermo, accuracy ± 0.2°C).

### High temperature tolerance assay

Carefully staged J3 worms from mixed stage plates incubated at 20°C were transferred singly to NGM plates, which were then shifted to 30°C. After seven days of incubation, the plates were inspected using a dissecting microscope and scored for their ability of the founder worm to give rise to fertile offspring. A strain was designated as Htt if the founder worm gave rise to two generations of offspring (the F2). A strain was designated Ltt if the founder was: killed by heat; arrested during development; developed to adulthood but was sterile; or gave rise only to F1 that were infertile. Three to 18 worms were tested and the mean number of individuals tested was five. For large-scale application of the Htt assay, a custom-made 30°C room (stability ± 0.5°C) was used to incubate the strains at the test temperature. Any strain that did not give reproducible results was eliminated from the final dataset, giving a final dataset of 289 strains.

### Statistical analysis

Statistical analysis was performed with RStudio software running R version 0.99.896 [49]. The data of the occurrence of Htt for all strains was fit with a generalised linear model (binomial distribution) using the glm command. The three variables for each strain were: the clade it belonged to; the altitude of its collections site; and the beetle species the strain was isolated from. See supplementary table 5 for the full data. Analysis of variance was performed using the anova command on the generalised linear model with the chi-square significance test. Bayesian Information Criterion analysis was performed using the bic command. The generalised linear model was fit to each variable independently and the output given as a weighted difference where the model that “best fits” the observed data has a value closest to 1. The results for the three models were 0.99 for altitude, 8.7×10^−11^ for clade and 3.0×10^−6^ for beetle.

### GWAS

GWAS was performed on the easyGWAS cloud server [35], where genomic sequence data and phenotype data were uploaded. The site allows for gene annotation to be automatically linked to the sequence that was uploaded. The beetle species that each strain was isolated from was included as a covariate, transformed as a dummy variable. A minor allele frequency filter was set to 10% and the EMMAX algorithm selected. Detailed, interactive results are available at https://easygwas.ethz.ch/data/public/datasets/.

### Data analysis for determining the altitudinal range for beetle species

For each sampling site visited, the number of beetles of each species collected was routinely noted for years 2010-2015. Also, the GPS coordinates and altitude of each site was noted. We define the altitudinal range for a beetle as being the range of altitudes where 95% of beetles were found and mean altitude of the collection site for all beetles of one species was also calculated.

### Construction of a phylogenetic tree

SNP data was extracted using a custom pipeline implemented in Perl and Unix as previously described [27]. Whole genome data for the stains used in the study was already available [25]. Variant bases were identified and the genotype of all strains at all variant positions was extracted from the genome sequence covered by all the contigs in *P. pacificus* draft genome hybrid assembly 1 [50]. 873932 SNPs were selected and concatenated. This sequence was used to construct a maximum likelihood tree using IQ-Tree [31]. The substitution rate model GTR+ASC was applied (where ASC corresponds to use of a SNP ascertainment bias model) and the tree was bootstrapped 1000 times using the ultrafast bootstrapping method [32].

### Inferred ancestral states

Bayesian analysis was performed using the ace (ancestral character estimation) command [51] implemented in the R phytools package [52]. A single rate model was applied and probabilities for either phenotype for each ancestor was calculated. The data was plotted with the R ggtree package with the probabilities at the internal nodes depicted as pie charts.

### Calculation of timeframe for adaptation

One generation is estimated to be between 3 days, being the time to develop from egg to fertile adult at 25°C, to 3 months, being the maximum time a dauer can survive. The number of generations it took to generate the diversity in clade C is estimated to be 5.6×10^5^ [37,38]. If one generation is 3 days, then 5.6×10^5^ generations = 3×5.6×10^5^ days or 1.68×10^6^ days. 365 days per year, then 1.68×10^6^ days = 4602 years. A similar logic gives the upper limit at 140000 years for a generation time of 3 months.

## Acknowledgments

We would like to thank: Jacques Rochat (Insectarium de Le Réunion), Matthias Hermann, Katie Morgan, Eduardo Moreno, and Jan Mayer for help sampling; Holger Brandl, Cameron Weadick, and Neel Prabh for help with statistical analysis; Waltraud Röseler and Andrea Zinke for strain maintenance; Heike Haussmann for freezing strains; Dominik Grimm for advice performing GWAS. M.L. was supported by an EMBO long-term fellowship (grant number ALTF:434-2010).

## Author contributions

ML and MK phenotyped strains. ML, AM and CR constructed the phylogenetic tree. ML and AM performed the GWAS. CR measured allele frequencies. ML, AM, CR, AAH and RJS conceived the study and wrote the paper.

## Abbreviations

Htt: high temperature tolerant
GWAS: genome wide association study
Aav: Anova analysis of variance
GE: Grand Etang
SB: Saint Benoit
SNPs: single nucleotide polymorphisms

---

**Supplementary figure 1.**
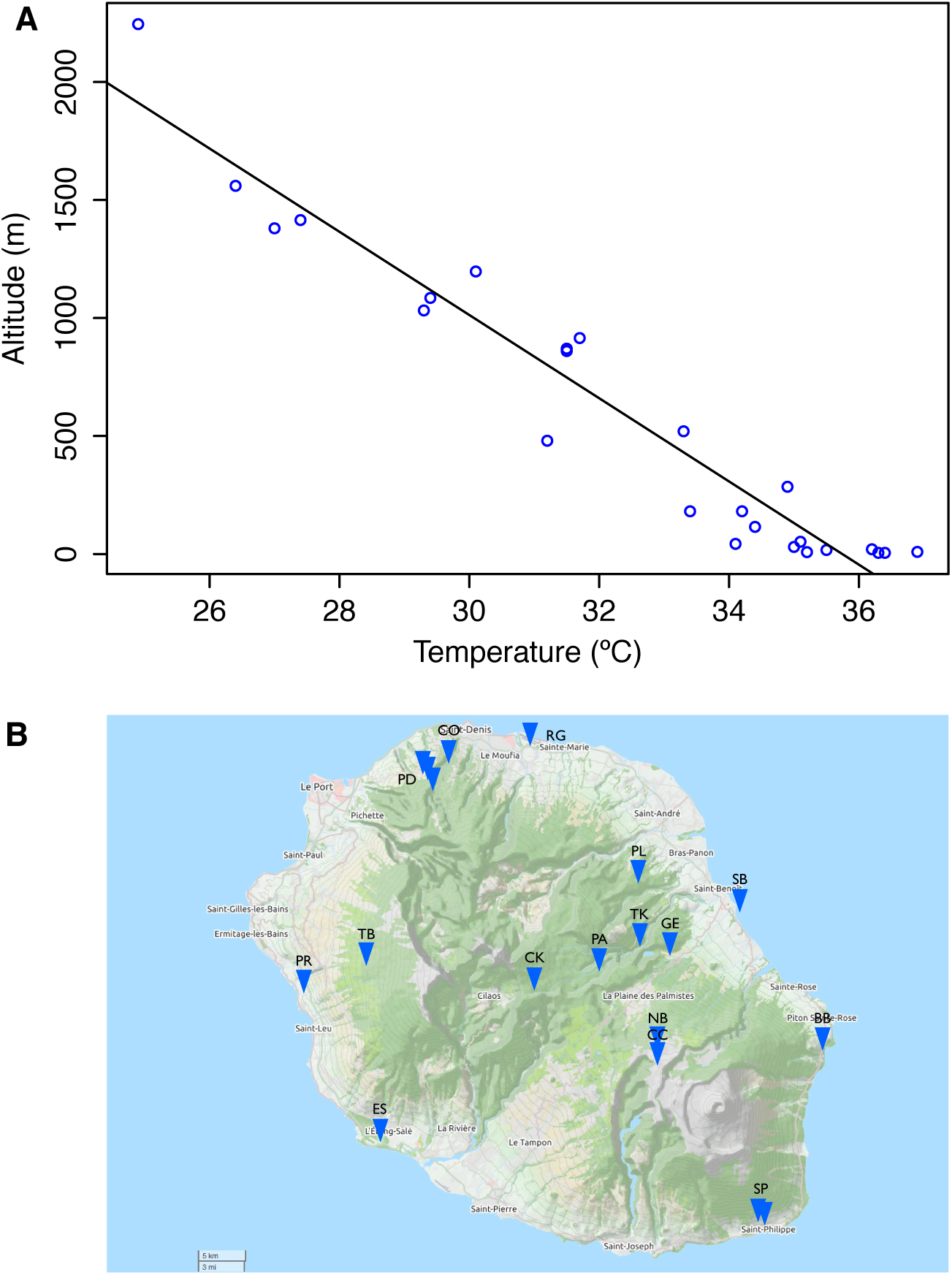
Altitude correlates with temperature on La Réunion. Panel A, the correlation between altitude and air temperature (adapted from Atlas Climatique de La Réunion). Panel B, locations where P. pacificus natural isolates were collected (see Table 1 for full location name and its altitude).

**Supplementary Figure 2.**
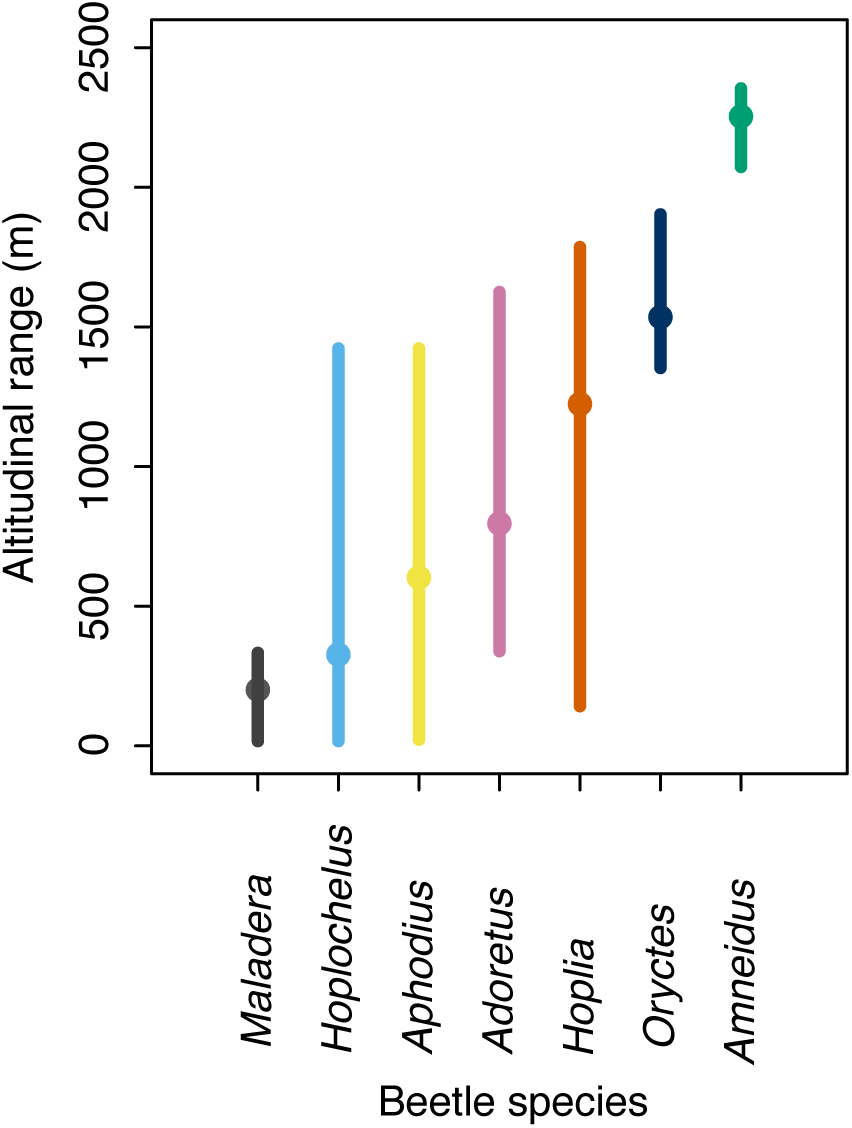
The altitudinal range of beetle species from La Réunion. The range of altitudes where beetle species are found. The vertical bar represents the maximum and minimum altitude each beetle species was found at. The point represents the mean altitude of the collection site for all beetles collected.

**Supplementary Figure 3.**
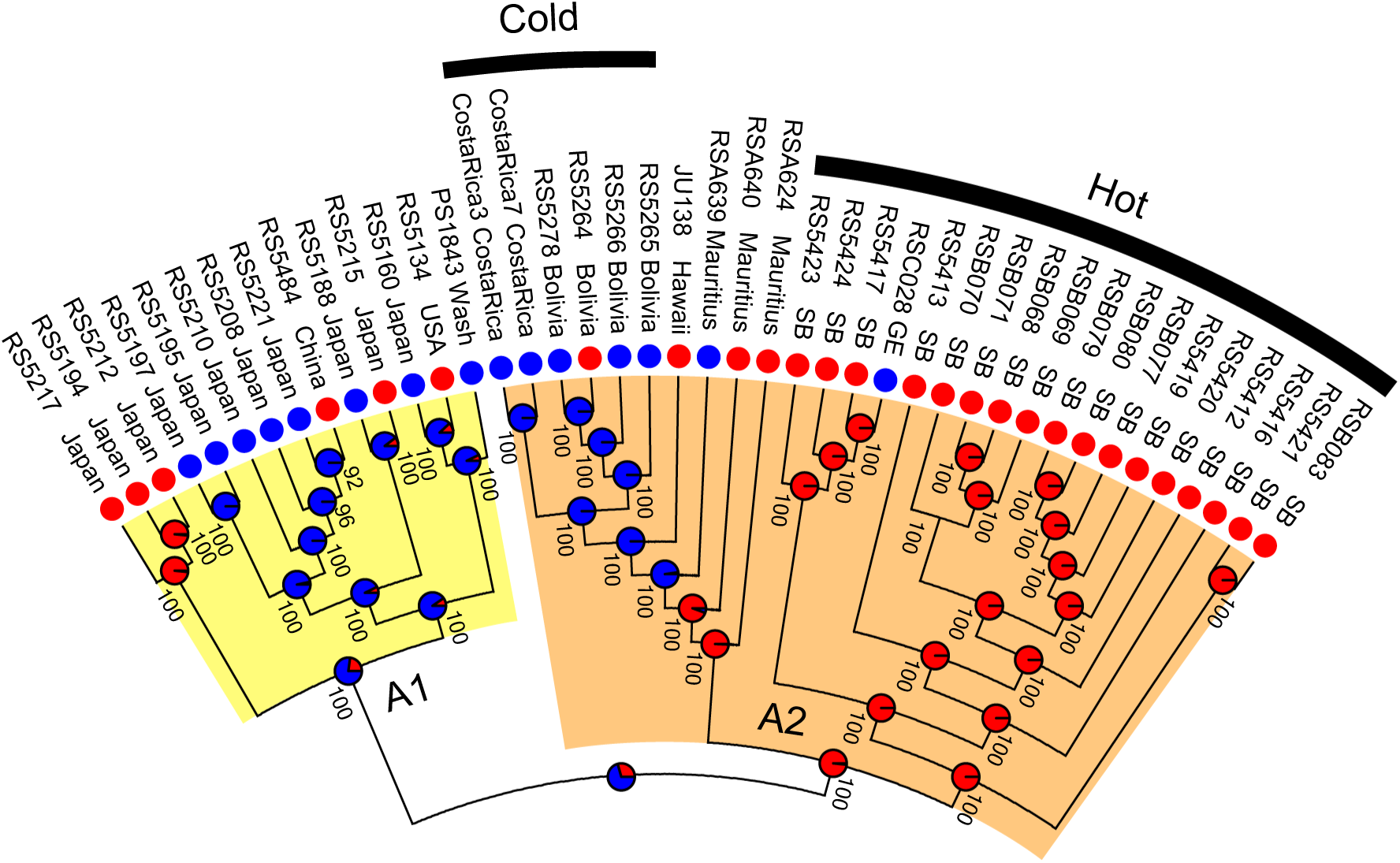
Cladogram of clade A1 and A2 strains showing parallel evolution of the trait. Cladogram based on clades A1 and A2 from the tree from figure 4. The phenotype of strains and the inferred ancestral phenotype are indicated by circles at the terminal nodes as in figure 4. Clade A1 is indicated by a yellow block and clade A2 by an orange block. The “hot” sub-clade is indicated by a black bar.

**Supplementary table 1.** The percentage of high temperature tolerant strains is higher at some low altitude collection sites. The number of strains collected for each location is shown as well as the number of strains able to give fertile offspring at 30°C (Htt strains) and the percentage of Htt strains per location (% Htt). Also shown are the total number of strains collected, the total number of Htt strains, and the percentage of Htt strains for the whole data set.

| Code | Name | Altitude | Strains | Htt Strains | % Htt |
| --- | --- | --- | --- | --- | --- |
| ES | Etang Salé | 17 | 3 | 1 | 33.3 |
| SB | Saint Benoit | 22 | 30 | 23 | 76.7 |
| RG | Airport Roland Garros | 26 | 1 | 0 | 0.0 |
| PR | Petite Ravine | 154 | 1 | 1 | 100.0 |
| BB2 | Bois Blanc 2 | 249 | 1 | 1 | 100.0 |
| BB3 | Bois Blanc 3 | 333 | 1 | 0 | 0.0 |
| SP09 | St Phillippe 9 | 339 | 34 | 11 | 32.4 |
| SP10 | St Phillippe 10 | 445 | 14 | 2 | 14.3 |
| SP11 | St Phillippe 11 | 511 | 24 | 8 | 33.3 |
| GE | Grand Etang | 527 | 11 | 0 | 0.0 |
| SP12 | St Phillippe 12 | 544 | 12 | 1 | 8.3 |
| PD2 | Plaine d'Affouches 2 | 691 | 12 | 5 | 41.7 |
| CO | Colorado | 694 | 41 | 7 | 17.1 |
| PL | Plaines des Lianes | 755 | 10 | 0 | 0.0 |
| PD3 | Plaine d'Affouches 3 | 809 | 6 | 0 | 0.0 |
| TK | Takamaka | 835 | 6 | 0 | 0.0 |
| PD4 | Plaine d'Affouches 4 | 886 | 9 | 2 | 22.2 |
| PA0 | Plaines des Palmiste 0 | 1199 | 1 | 0 | 0.0 |
| SS | Sans Soucie | 1353 | 10 | 0 | 0.0 |
| TB | Trois Bassin | 1413 | 30 | 0 | 0.0 |
| PA1 | Plaines des Palmiste 1 | 1466 | 2 | 0 | 0.0 |
| PA3 | Plaines des Palmiste 3 | 1627 | 3 | 0 | 0.0 |
| NB | Nez de Bouef | 2146 | 7 | 0 | 0.0 |
| CK | Coteau Kerveguen | 2150 | 10 | 0 | 0.0 |
| CC | Cratere commerson | 2327 | 10 | 0 | 0.0 |
|  |  | sum | 289 | 62 | 21.5 |

**Supplementary table 2.**
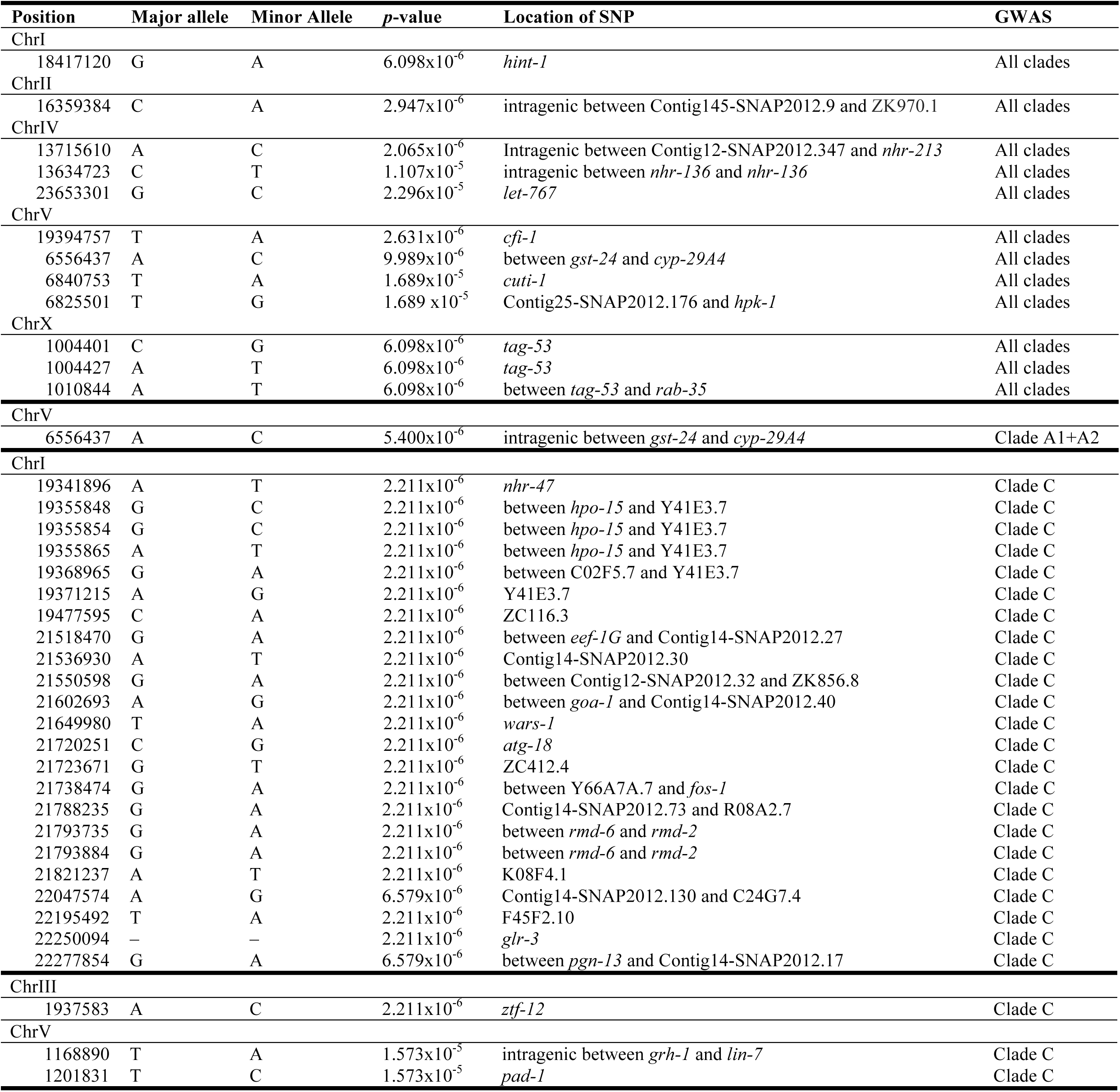
SNPs identified by GWAS.

| Position | Major allele | Minor Allele | p-value | Location of SNP | GWAS |
| --- | --- | --- | --- | --- | --- |
| ChrI |  |  |  |  |  |
| 18417120 | G | A | 6.098x10<sup>-6</sup> | *hint-1* | All clades |
| ChrII |  |  |  |  |  |
| 16359384 | C | A | 2.947x10<sup>-6</sup> | intragenic between Contig145-SNAP2012.9 and ZK970.1 | All clades |
| ChrIV |  |  |  |  |  |
| 13715610 | A | C | 2.065x10<sup>-6</sup> | Intragenic between Contig12-SNAP2012.347 and *nhr-213* | All clades |
| 13634723 | C | T | 1.107x10<sup>-5</sup> | intragenic between *nhr-136* and *nhr-136* | All clades |
| 23653301 | G | C | 2.296x10<sup>-5</sup> | *let-767* | All clades |
| ChrV |  |  |  |  |  |
| 19394757 | T | A | 2.631x10<sup>-6</sup> | *cfi-1* | All clades |
| 6556437 | A | C | 9.989x10<sup>-6</sup> | between *gst-24* and *cyp-29A4* | All clades |
| 6840753 | T | A | 1.689x10<sup>-5</sup> | *cuti-1* | All clades |
| 6825501 | T | G | 1.689 x10<sup>-5</sup> | Contig25-SNAP2012.176 and *hpk-1* | All clades |
| ChrX |  |  |  |  |  |
| 1004401 | C | G | 6.098x10<sup>-6</sup> | *tag-53* | All clades |
| 1004427 | A | T | 6.098x10<sup>-6</sup> | *tag-53* | All clades |
| 1010844 | A | T | 6.098x10<sup>-6</sup> | between *tag-53* and *rab-35* | All clades |
| ChrV |  |  |  |  |  |
| 6556437 | A | C | 5.400x10<sup>-6</sup> | intragenic between *gst-24* and *cyp-29A4* | Clade A1+A2 |
| ChrI |  |  |  |  |  |
| 19341896 | A | T | 2.211x10<sup>-6</sup> | *nhr-47* | Clade C |
| 19355848 | G | C | 2.211x10<sup>-6</sup> | between *hpo-15* and Y41E3.7 | Clade C |
| 19355854 | G | C | 2.211x10<sup>-6</sup> | between *hpo-15* and Y41E3.7 | Clade C |
| 19355865 | A | T | 2.211x10<sup>-6</sup> | between *hpo-15* and Y41E3.7 | Clade C |
| 19368965 | G | A | 2.211x10<sup>-6</sup> | between C02F5.7 and Y41E3.7 | Clade C |
| 19371215 | A | G | 2.211x10<sup>-6</sup> | Y41E3.7 | Clade C |
| 19477595 | C | A | 2.211x10<sup>-6</sup> | ZC116.3 | Clade C |
| 21518470 | G | A | 2.211x10<sup>-6</sup> | between *eef-1G* and Contig14-SNAP2012.27 | Clade C |
| 21536930 | A | T | 2.211x10<sup>-6</sup> | Contig14-SNAP2012.30 | Clade C |
| 21550598 | G | A | 2.211x10<sup>-6</sup> | between Contig12-SNAP2012.32 and ZK856.8 | Clade C |
| 21602693 | A | G | 2.211x10<sup>-6</sup> | between *goa-1* and Contig14-SNAP2012.40 | Clade C |
| 21649980 | T | A | 2.211x10<sup>-6</sup> | *wars-1* | Clade C |
| 21720251 | C | G | 2.211x10<sup>-6</sup> | *atg-18* | Clade C |
| 21723671 | G | T | 2.211x10<sup>-6</sup> | ZC412.4 | Clade C |
| 21738474 | G | A | 2.211x10<sup>-6</sup> | between Y66A7A.7 and *fos-1* | Clade C |
| 21788235 | G | A | $2.211 \times 10^{-6}$ | Contig14-SNAP2012.73 and R08A2.7 | Clade C |
| 21793735 | G | A | $2.211 \times 10^{-6}$ | between *rmd-6* and *rmd-2* | Clade C |
| 21793884 | G | A | $2.211 \times 10^{-6}$ | between *rmd-6* and *rmd-2* | Clade C |
| 21821237 | A | T | $2.211 \times 10^{-6}$ | K08F4.1 | Clade C |
| 22047574 | A | G | $6.579 \times 10^{-6}$ | Contig14-SNAP2012.130 and C24G7.4 | Clade C |
| 22195492 | T | A | $2.211 \times 10^{-6}$ | F45F2.10 | Clade C |
| 22250094 | – | – | $2.211 \times 10^{-6}$ | *glr-3* | Clade C |
| 22277854 | G | A | $6.579 \times 10^{-6}$ | between *pgn-13* and Contig14-SNAP2012.17 | Clade C |
| ChrIII |  |  |  |  |  |
| 1937583 | A | C | $2.211 \times 10^{-6}$ | *ztf-12* | Clade C |
| ChrV |  |  |  |  |  |
| 1168890 | T | A | $1.573 \times 10^{-5}$ | intragenic between *grh-1* and *lin-7* | Clade C |
| 1201831 | T | C | $1.573 \times 10^{-5}$ | *pad-1* | Clade C |

**Supplementary table 3.** Allele frequencies of GWAS hits show some alleles are variable across clades while other alleles unique to one clade. For each position that was identified as being associated with the Htt phenotype, the allele frequency was calculated for all stains in each clade. SNPs that are variable in many clades are highlighted in yellow, those that are variable in only one clade are highlighted in orange, red and blue corresponding to clades A1, A2 and C respectively.

| SNP position | GWAS | Allele frequencies in individual clades |  |  |  |  |  |
| --- | --- | --- | --- | --- | --- | --- | --- |
|  |  | A1 | A2 | A3 | B | C | D |
| ChrI:18417120 | All clades | G:0.79 A:0.21 | A:1.00 | G:1.00 | A:1.00 | G:1.00 | G:1.00 |
| ChrII:16359384 | All clades | C:1.00 | C:0.29 A:0.71 | C:1.00 | A:1.00 | C:1.00 | C:1.00 |
| ChrIV:13634723 | All clades | C:0.29 T:0.71 | C:1.00 | C:1.00 | C:1.00 | T:0.01 C:0.99 | T:1.00 |
| ChrIV:13715610 | All clades | A:0.71 C:0.29 | C:1.00 | C:1.00 | A:1.00 | A:1.00 | A:1.00 |
| ChrIV:23653301 | All clades | C:0.21 G:0.79 | C:1.00 | C:0.80 G:0.20 | G:1.00 | G:1.00 | G:1.00 |
| ChrV:19394757 | All clades | A:0.64 T:0.36 | T:0.68 A:0.32 | T:1.00 | A:1.00 | T:1.00 | T:1.00 |
| ChrV:6825501 | All clades | T:0.71 G:0.29 | G:1.00 | T:0.13 G:0.87 | T:1.00 | T:1.00 | T:1.00 |
| ChrV:6840753 | All clades | T:0.71 A:0.29 | A:1.00 | T:0.13 A:0.87 | T:1.00 | T:1.00 | T:1.00 |
| ChrX:1004401 | All clades | C:0.79 G:0.21 | G:1.00 | C:1.00 | G:1.00 | C:1.00 | C:1.00 |
| ChrX:1004427 | All clades | T:0.21 A:0.79 | T:1.00 | A:1.00 | T:1.00 | A:1.00 | A:1.00 |
| ChrX:1010844 | All clades | T:0.21 A:0.79 | T:1.00 | A:1.00 | T:1.00 | A:1.00 | A:1.00 |
| ChrV:6556437 | Clade A1+A2 | C:0.64 A:0.36 | C:0.29 A:0.71 | A:1.00 | A:1.00 | A:0.94 C:0.06 | A:0.19 C:0.81 |
| ChrI:19341896 | Clade C | A:1.00 | A:1.00 | A:1.00 | A:1.00 | T:0.29 A:0.71 | A:1.00 |
| ChrI:19355848 | Clade C | C:1.00 | C:1.00 | C:1.00 | C:1.00 | C:0.71 G:0.29 | C:1.00 |
| ChrI:19355854 | Clade C | G:1.00 | G:1.00 | G:1.00 | G:1.00 | G:0.71 C:0.29 | G:1.00 |
| ChrI:19355865 | Clade C | A:1.00 | A:1.00 | A:1.00 | A:1.00 | T:0.29 A:0.71 | A:1.00 |
| ChrI:19368965 | Clade C | A:1.00 | A:1.00 | A:1.00 | A:1.00 | A:0.71 G:0.29 | A:1.00 |
| ChrI:19371215 | Clade C | A:1.00 | A:1.00 | A:1.00 | A:1.00 | G:0.29 A:0.71 | A:1.00 |
| ChrI:19477595 | Clade C | A:1.00 | A:1.00 | A:1.00 | A:1.00 | C:0.71 A:0.29 | A:1.00 |
| ChrI:21518470 | Clade C | G:1.00 | G:1.00 | G:1.00 | G:1.00 | G:0.71 A:0.29 | G:1.00 |
| ChrI:21536930 | Clade C | A:1.00 | A:1.00 | A:1.00 | A:1.00 | T:0.29 A:0.71 | A:1.00 |
| ChrI:21550598 | Clade C | A:1.00 | A:1.00 | A:1.00 | A:1.00 | G:0.29 A:0.71 | A:1.00 |
| ChrI:21602693 | Clade C | A:1.00 | A:1.00 | A:1.00 | A:1.00 | G:0.29 A:0.71 | A:1.00 |
| ChrI:21649980 | Clade C | T:1.00 | T:1.00 | T:1.00 | T:1.00 | T:0.71 A:0.29 | T:1.00 |
| ChrI:21720251 | Clade C | C:1.00 | C:1.00 | C:1.00 | C:1.00 | G:0.29 C:0.71 | C:1.00 |
| ChrI:21723671 | Clade C | G:1.00 | G:1.00 | G:1.00 | G:1.00 | G:0.71 T:0.29 | G:1.00 |
| ChrI:21738474 | Clade C | G:1.00 | G:1.00 | G:1.00 | G:1.00 | G:0.71 A:0.29 | G:1.00 |
| ChrI:21788235 | Clade C | G:1.00 | G:1.00 | G:1.00 | G:1.00 | G:0.71 A:0.29 | G:1.00 |
| ChrI:21793735 | Clade C | G:1.00 | G:1.00 | G:1.00 | G:1.00 | A:0.29 G:0.71 | G:1.00 |
| ChrI:21793884 | Clade C | G:1.00 | G:1.00 | G:1.00 | G:1.00 | G:0.71 A:0.29 | G:1.00 |
| ChrI:21821237 | Clade C | A:1.00 | A:1.00 | A:1.00 | A:1.00 | A:0.71 T:0.29 | A:1.00 |
| ChrI:22047574 | Clade C | A:1.00 | A:1.00 | A:1.00 | A:1.00 | G:0.28 A:0.72 | A:1.00 |
| ChrI:22195492 | Clade C | T:1.00 | T:1.00 | T:1.00 | T:1.00 | A:0.29 T:0.71 | T:1.00 |
| ChrI:22250094 | Clade C | C:1.00 | C:1.00 | C:1.00 | C:1.00 | T:0.29 C:0.71 | C:1.00 |
| ChrI:22277854 | Clade C | G:1.00 | G:1.00 | G:1.00 | G:1.00 | A:0.28 G:0.72 | G:1.00 |
| ChrIII:1937583 | Clade C | C:1.00 | C:1.00 | C:1.00 | C:1.00 | C:0.29 A:0.71 | C:1.00 |
| ChrV:1168890 | Clade C | T:1.00 | A:0.14 T:0.86 | T:1.00 | T:1.00 | T:0.81 A:0.19 | T:0.95 A:0.05 |
| ChrV:1201831 | Clade C | T:1.00 | T:0.86 C:0.14 | T:1.00 | T:1.00 | T:0.81 C:0.19 | T:0.95 C:0.05 |

**Supplementary table 4.** Genes containing or near to SNPs identified by GWAS. Predicted gene function based of protein family or closest homolog in *C. elegans*.

| gene | protein | function | References |
| --- | --- | --- | --- |
| *atg-18* | Autophagy | essential for normal dauer morphogenesis and autophagy in daf-2 mutant animals | Wormbase |
| C24G7.4 | ACD-2 | sodium channel involved in mechanosensation | Wormbase |
| *cfi-1* | CEM Fate inhibitor | Cell fate determination of sensory neurons | Wormbase |
| *cuti-1* | cuticle and epithelial Integrity | cuticle development, embryo development, locomotion, the molting cycle, nematode larval development and reproduction | Wormbase |
| *cyp-29A4* | cytochrome P450 | electron transport, xenobiotic detoxification, fatty acid synthesis | Wormbase |
| *eef-1G* | Eukaryotic translation Elongation Factor | involved in embryo development and reproduction. RNAi gives reduced broodsize, gonad defects and slow growth | Wormbase |
| F45F2.10 | ankyrin repeat domain 36 | involved in embryo development and reproduction: RNAi gives increased lifespan in dauers, embryonic lethal, maternal sterile, and sterile progeny | Wormbase |
| *fos-1* | B-Zip transcription factor | anchor cell invasion of the vulval epithelium and proper vulval and uterine development, fertility, and oogenesis | Wormbase |
| *goa-1* | G protein, O, Alpha subunit | regulation of locomotion, egg-laying, male mating, and olfactory-mediated behaviors; also required for asymmetric cell division in the early embryo | Wormbase |
| *glb-10* | globin | chemosensation | Wormbase |
| *grh-1* | Transcription factor | Cuticle synthesis and embryonic development | Wormbase |
| *glr-3* | GLutamate Receptor family |  | Wormbase |
| *gst-24* | glutathione S-transferase | Oxidative stress response | Wormbase |
| *hint-1* | histidine triad nucleotide binding protein 1 |  | Wormbase |
| *hpk-1* | Homeodomain interacting Protein Kinase | localized to the nucleus and is expressed in cleavage stage embryos | Wormbase |
| *hpo-15* | Hypersensitive to PORE-forming toxin | predicted oxidoreductase activity | Wormbase |
| K08F4.1 | chromosome transmission fidelity factor 18 | cellular response to DNA damage stimulus | Wormbase |
| *let-767* | lethal | required for embryogenesis, molting, and female reproduction | Wormbase |
| *lin-7* | abnormal cell LINEage | Required for cell polarity in vulva precursor cells | Wormbase |
| M03B6.2 | MCT-3 | Solute carriers | Wormbase |
| *mua-3* | Vitrin | apoptotic process, cell-matrix adhesion, locomotion, nematode larval development and reproduction | Wormbase |
| *nhr-47* | nuclear hormone receptor | Expressed in response to estradiol exposure | Wormbase |
| *nhr-213* | nuclear hormone receptor | memory | Wormbase |
| *nhr-136* | nuclear hormone receptor | Starvation response | Wormbase |
| *pad-1* | PAtternning Defective | Required for embryonic development | Wormbase |
| *pgn-13* |  |  |  |
| *rab-35* | Rab GTPase | intracellular vesicular trafficking and for regulation of endo- and exocytosis | Wormbase |
| *rmd-2* | Regulator of Microtubule Dynamics | involved in striated muscle myosin thick filament assembly | Wormbase |
| *rmd-6* | Regulator of Microtubule Dynamics | RNAi gives phenotypes in endosome recycling lysosome morphology defects | Wormbase |
| R08A2.7 |  |  |  |
| *tag-35* |  |  |  |
| *wars-1* | tryptophanyl Amino-acyl tRNA Synthetase | affects fertility, embryonic viability, growth, and locomotion | Wormbase |
| Y66A7A.7 |  |  |  |
| *ztf-12* | SUP-37 | involved in embryo development, embryonic digestive tract morphogenesis, nematode larval development and reproduction | Wormbase |
| ZC116.3 | CUBILIN |  | Wormbase |
| ZC412.4 |  |  |  |
| ZK856.8 | Chpf-1 | involved in locomotion | Wormbase |
| Y41E3.7 |  |  |  |
| ZK970.1 | NEP-26 | A zinc metallopeptidases, found on the outer surface of animal cells that negatively regulate small signaling peptides | Wormbase |

**Supplementary Table 5.**
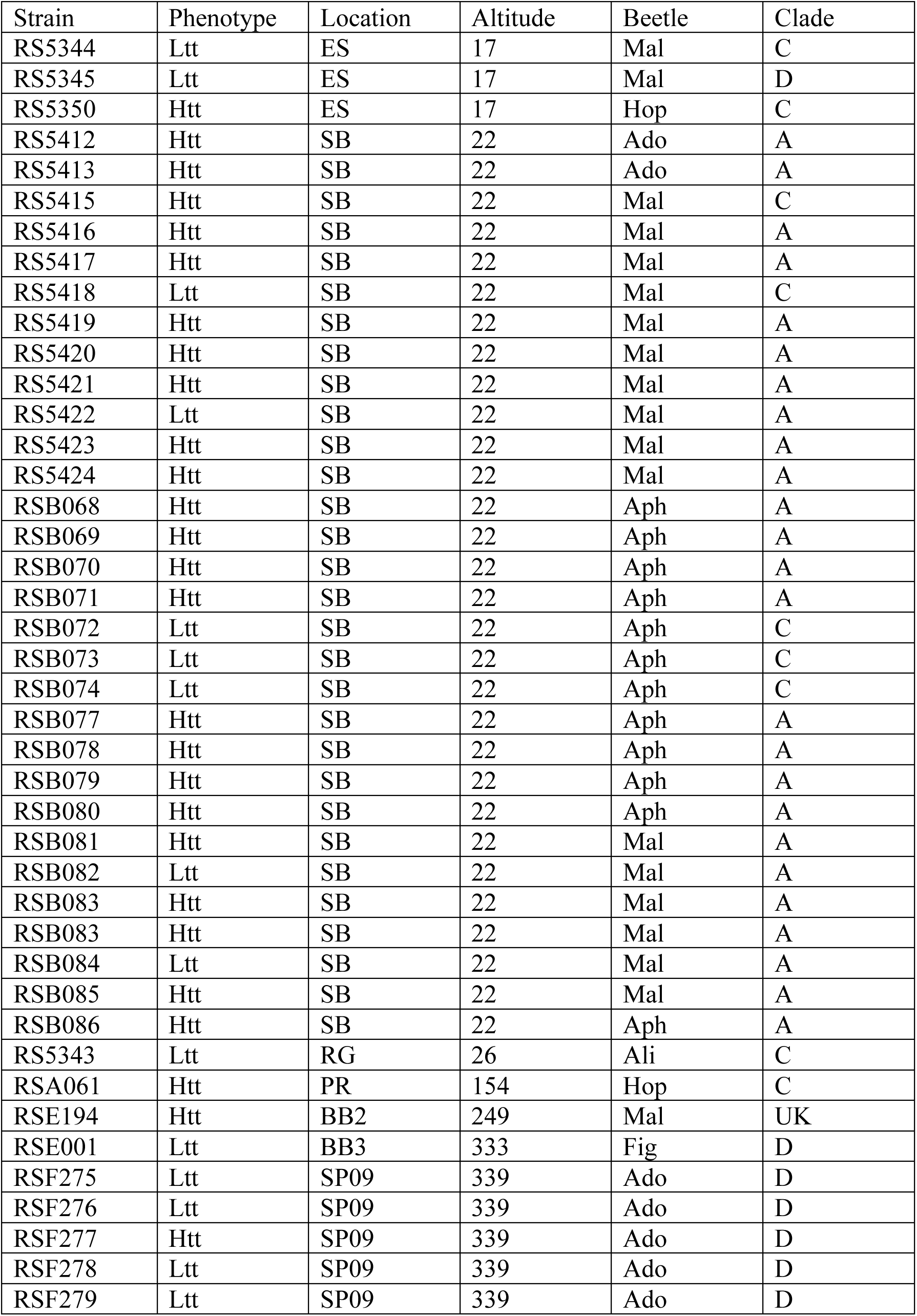
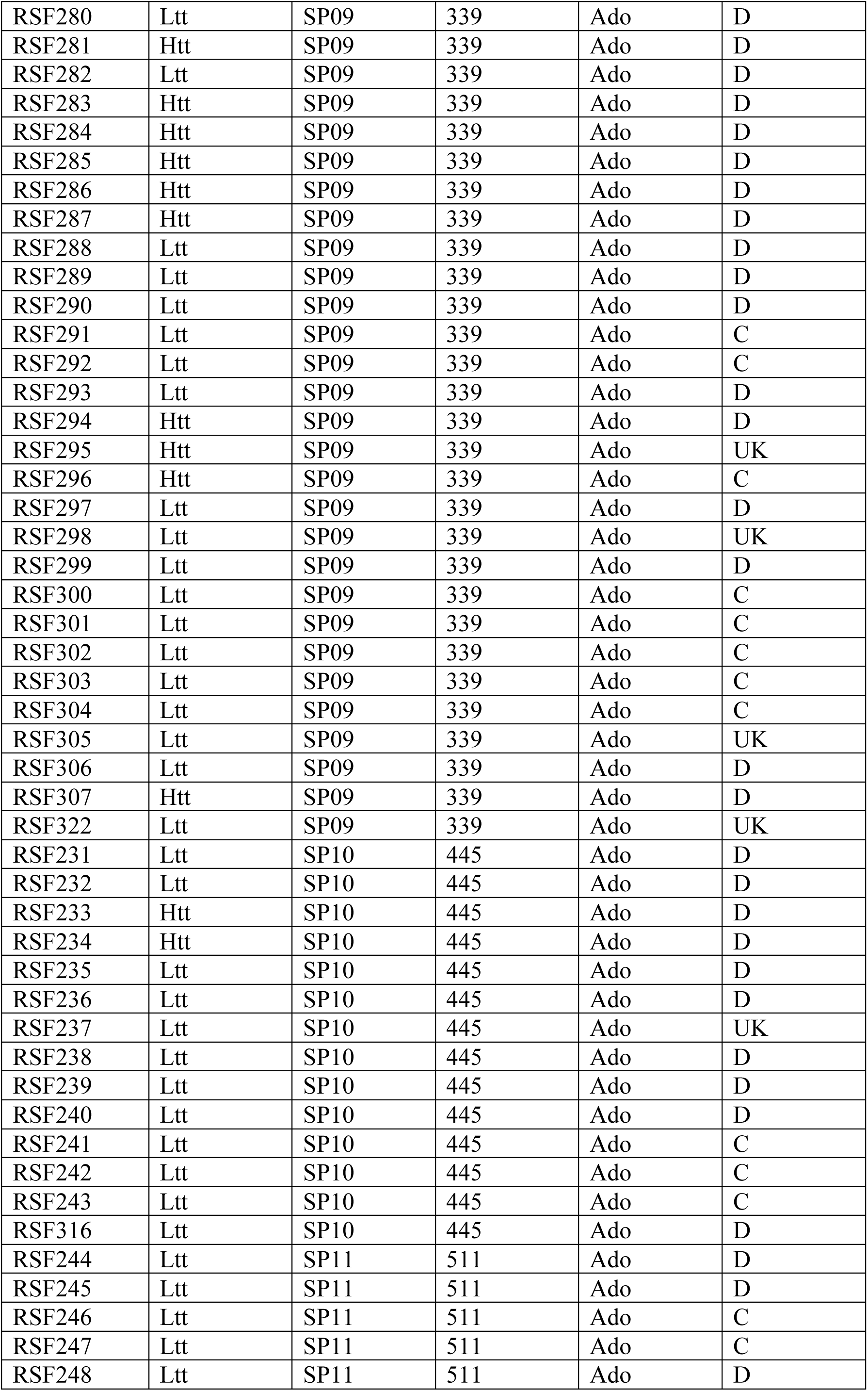
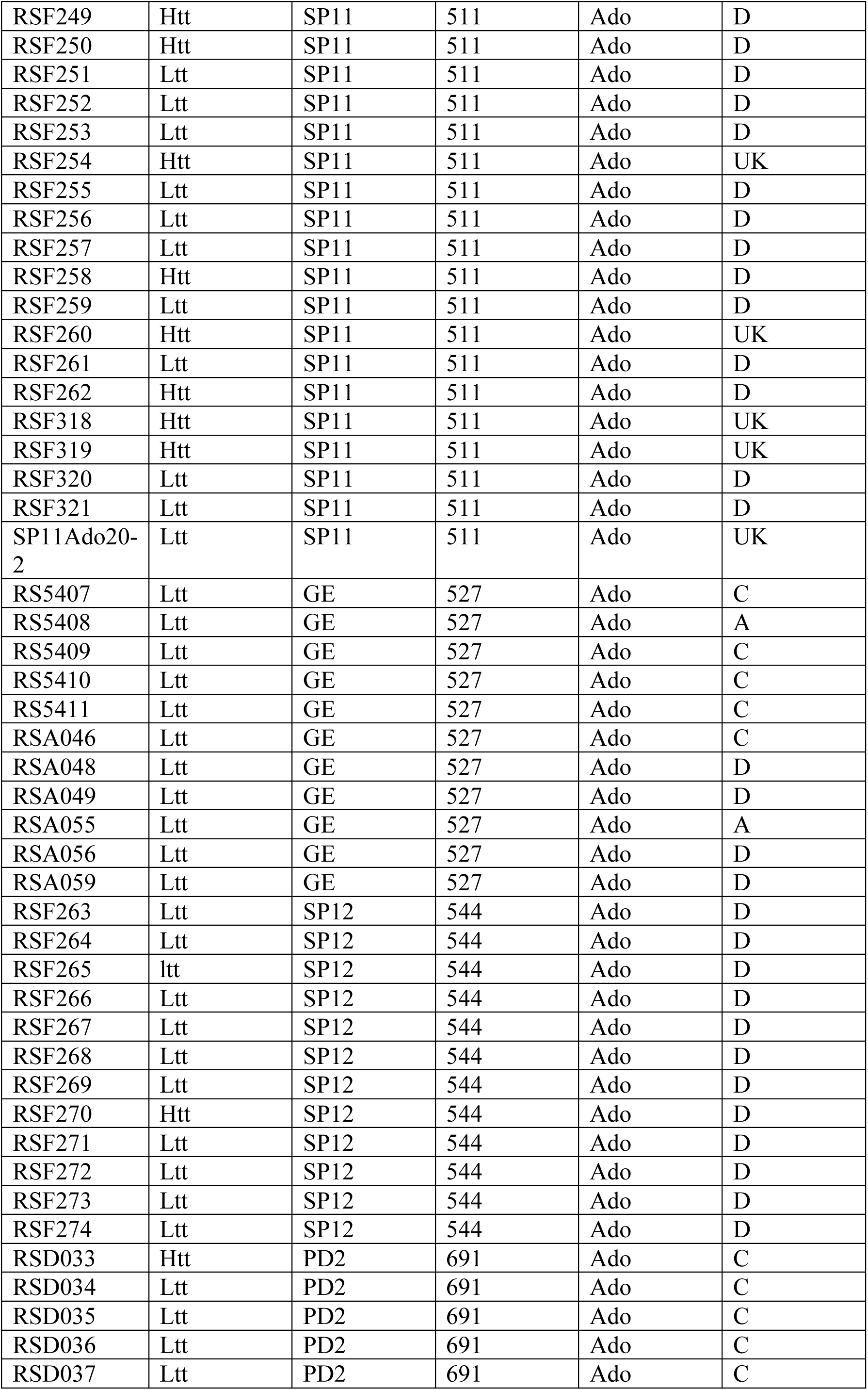
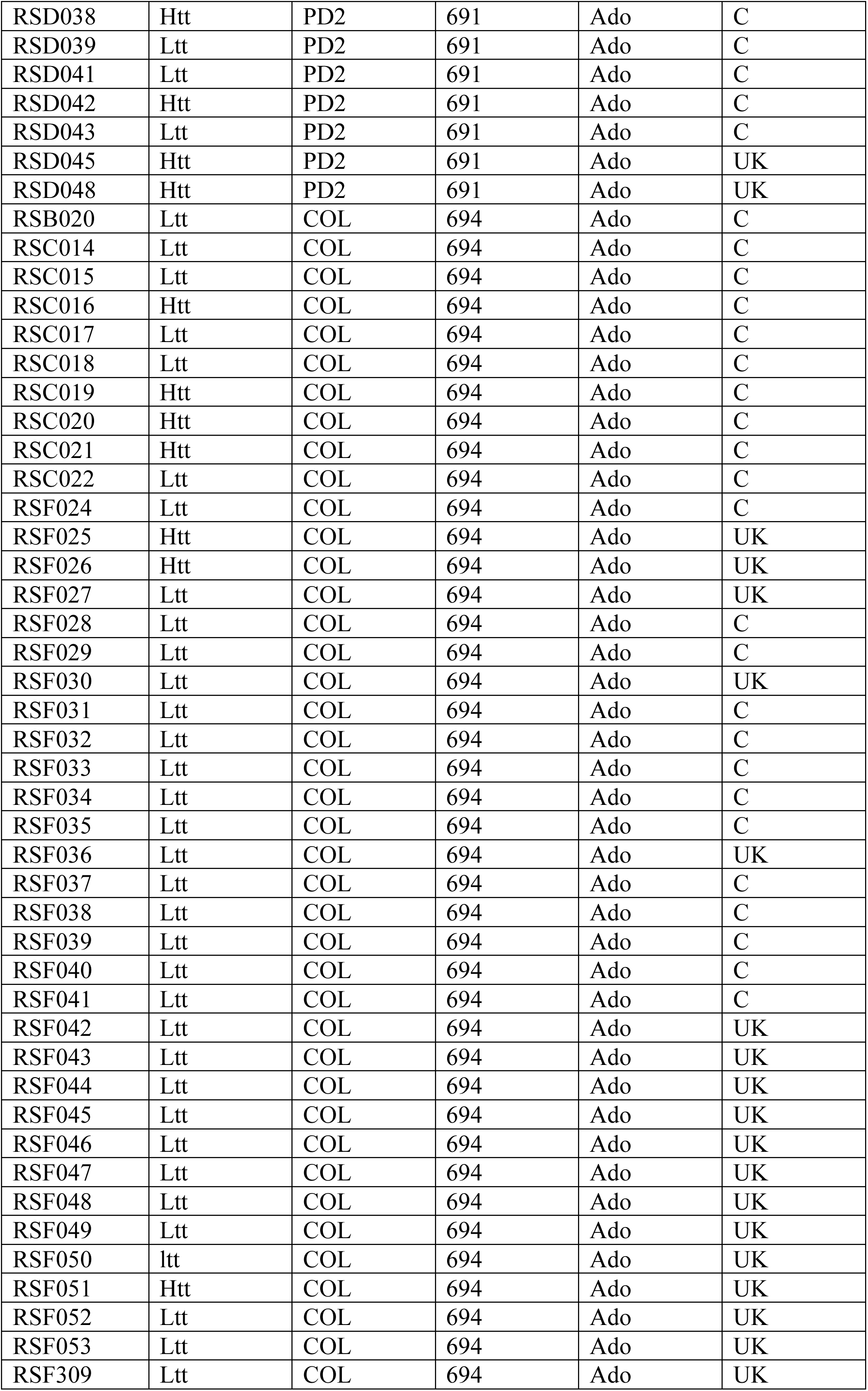
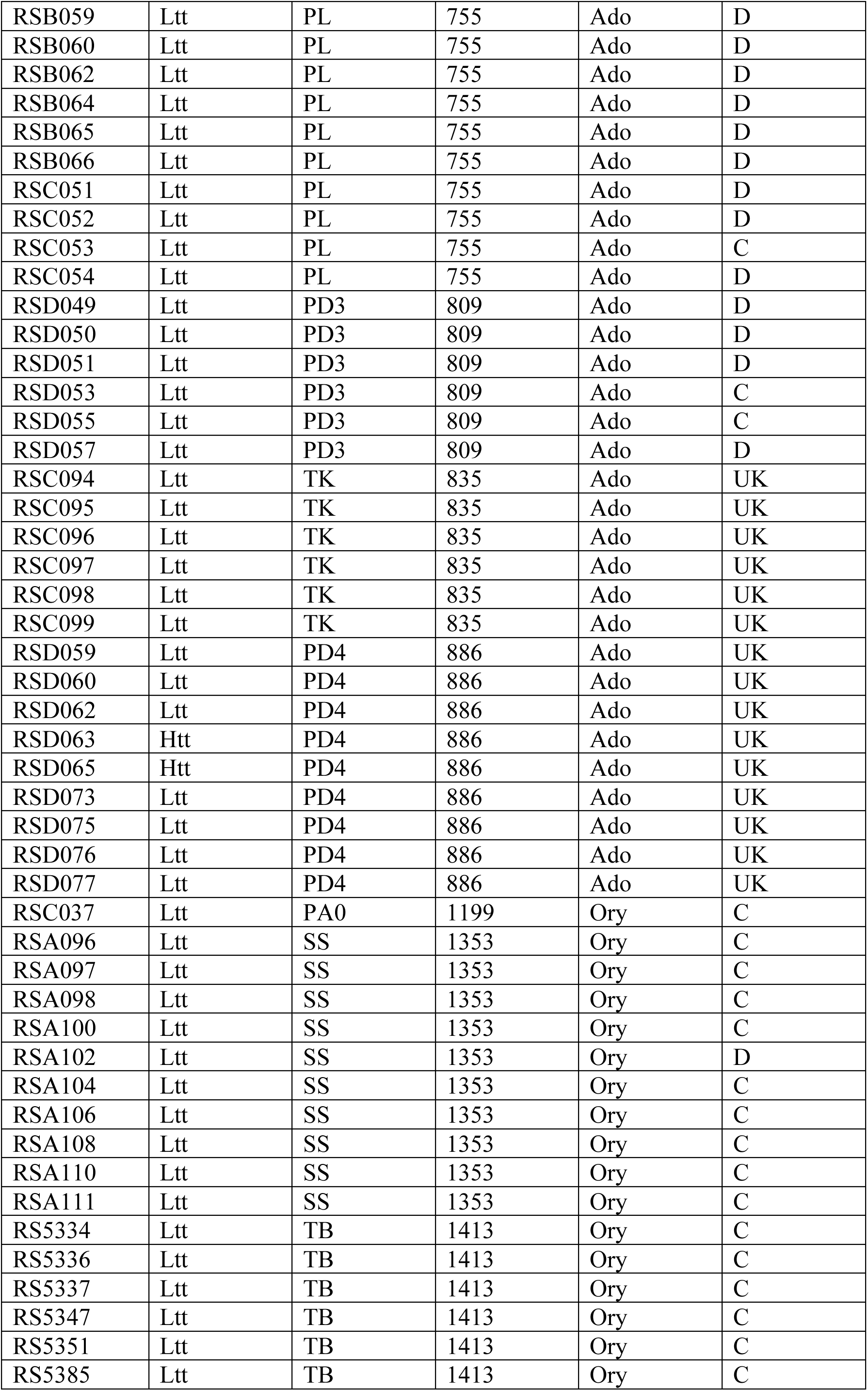
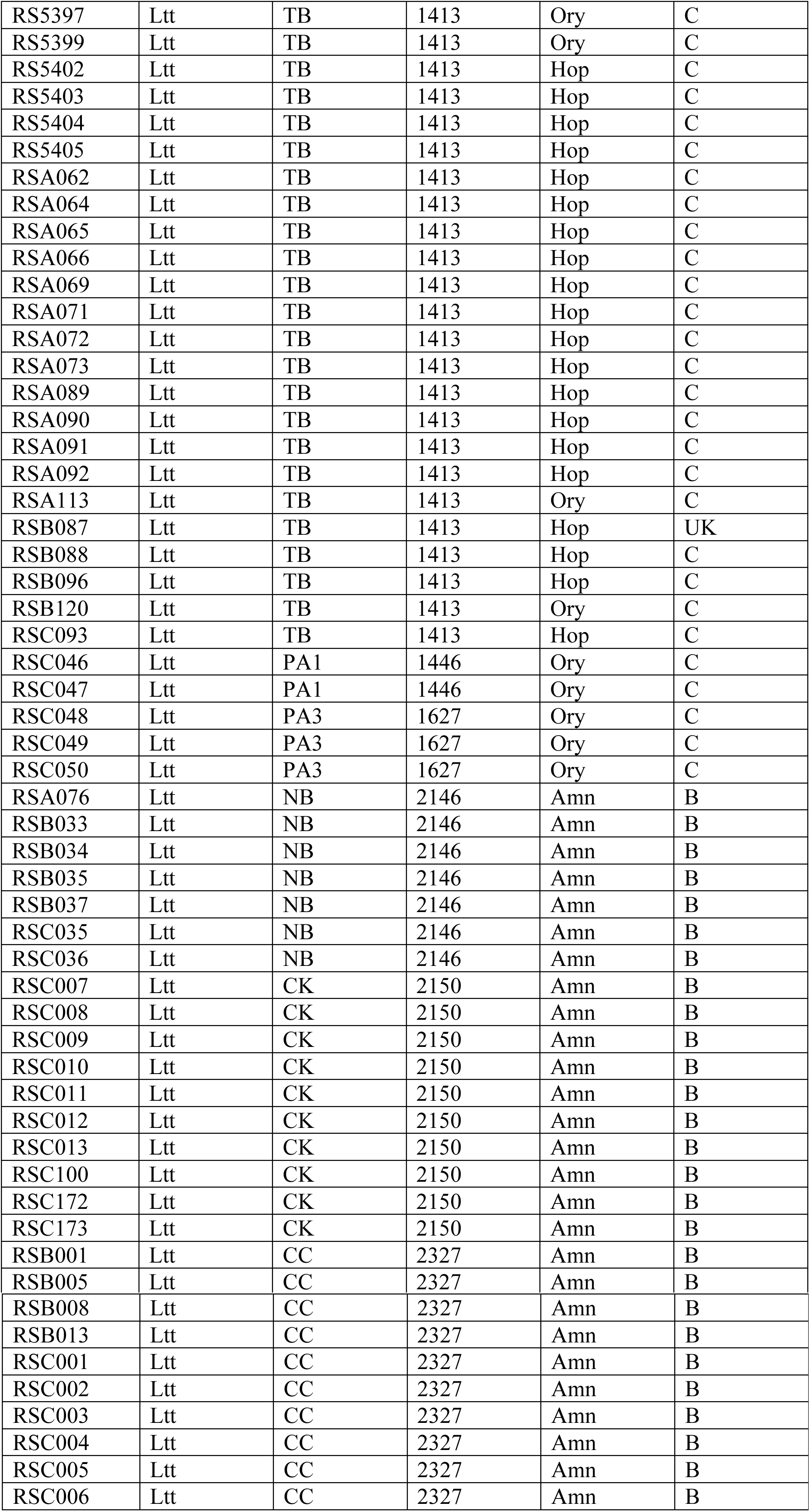
List of strains used in the study. The table lists for each strain its phenotype, it collection site, its beetle host and the mitochondrial clade to which it belongs. For strain that were not sequenced, no clade information is available, indicated by UK. See table 1 for the full name of the collection site. The altitude in meters for each site is also noted.

